# From Breath to Behavior: Respiratory Features Predict Visual Detection Performance

**DOI:** 10.64898/2026.03.18.712638

**Authors:** Emily Skog, Deepa Issar, Madison Grigg, Samantha E. Nelson, Jana M. Kainerstorfer, Matthew A. Smith

## Abstract

Breathing is a continuous bodily rhythm that not only sustains physiology but also shapes brain function and behavior. Here we investigated how respiration interacts with perceptual performance in nonhuman primates performing a visual detection task. Using continuous recordings, we extracted detailed features from each respiratory cycle including timing, duration, phase, depth, and volume, aligned to trial onset. Analyses revealed that timing-related features, such as inhalation onset and the respiration length, were the most reliable markers of trial outcome, whereas amplitude-based measures contributed less consistently. These findings demonstrate that the temporal structure of breathing, rather than its magnitude, plays a dominant role in shaping behavior on a moment-to-moment basis. By uncovering how fine-grained features of respiration align with perceptual success, our work highlights respiration as a strong correlate of cognition and highlights the value of feature-based approaches for linking interoceptive rhythms to behavior.

## INTRODUCTION

Respiration is a continuous, partially volitional internal bodily rhythm that has functions that extend far beyond oxygen metabolism. Accumulating evidence from human and animal studies suggests that respiration dynamically interacts with neural circuits and modulates cognitive performance. For example, the phase, volume and rate of breathing have been shown to entrain neural oscillations across multiple brain regions, including the prefrontal cortex^1–4^, hippocampus^1,2,5^, and sensory areas^2,6–9^. These respiration-locked fluctuations similarly influence perceptual capabilities, which were reflected in changes in reaction times^10–12^ and task accuracy^2,13–17^. Thus, the act of breathing is tightly interwoven with how the brain perceives, decides, and acts.

More broadly, respiration is a central component of interoception, the process by which the brain senses and integrates internal bodily signals^18^. Interoception is fundamental for maintaining physiological balance and guiding adaptive behavior, and its disruption has been linked to neuropsychiatric and neurodegenerative disorders, including schizophrenia^19–21^, depression^22–24^, and Alzheimer’s disease^25,26^. Studying respiration therefore offers a unique lens into how interoceptive signals modulate neural activity and cognition.

Prior research on respiration and behavior has largely emphasized slower, minutes-to-hours timescales^27,28^, such as in sleep^29,30^, anxiety^31–33^, or respiratory disease detection^34–36^. Far less is known about the interactions between respiration and behavior at finer resolutions, such as moment-to-moment perceptual decisions. In addition, many analyses rely on single-dimensional summaries of breathing (e.g., rate, depth, or a single phase value)^14,15^, which capture broad dynamics but may overlook the rich temporal and structural features embedded in the respiratory signal. This is particularly important because respiration is inherently variable. Features such as phase, depth, flow, duration, and rate covary in complex, nonlinear ways from cycle to cycle. This variability introduces a rich but underexamined source of information^27,37^. Additionally, general studies often emphasize that neural and behavioral measures are aligned to inhalation, treating it as a key respiratory feature for task success, yet this may obscure subtler features distributed across the entire respiratory cycle^15,38^.

To overcome these limitations, we present a novel feature-based framework that decodes behavior directly from trial-level respiratory dynamics in a perceptual decision-making task in non-human primates (NHPs). Using high-resolution respiration recordings from NHPs primates performing a visual detection task, we extracted a comprehensive set of features from each respiratory cycle, including timing, phase, duration, depth, and volume, aligned to the onset of each trial. We then used these features to predict behavioral outcomes on a single-trial basis, using interpretable machine learning models. We found that timing-related features (such as rate) were consistently more predictive of behavioral performance than amplitude-based features (such as depth or volume). Overall, this study supports a role for respiration in modulating behavior.

## RESULTS

### Respiration was recorded during a visual perceptual task in NHPs

We recorded high-resolution respiration waveforms from two male rhesus macaque monkeys (Monkey RA and Monkey AB) during a visual color-change detection task (Figure 1A). Monkeys fixated on a central dot and withheld eye movements until the fixation changed color. Then they made a saccade to a peripheral target to obtain reward (Figure 1B). Trials were classified as correct, false alarm, or miss, as described in the STAR Methods. Perceptual difficulty was manipulated by varying the magnitude of the color change (Figure 1C). This design allowed us to examine how respiration interacts with varying task difficulty.

**Figure 1.**
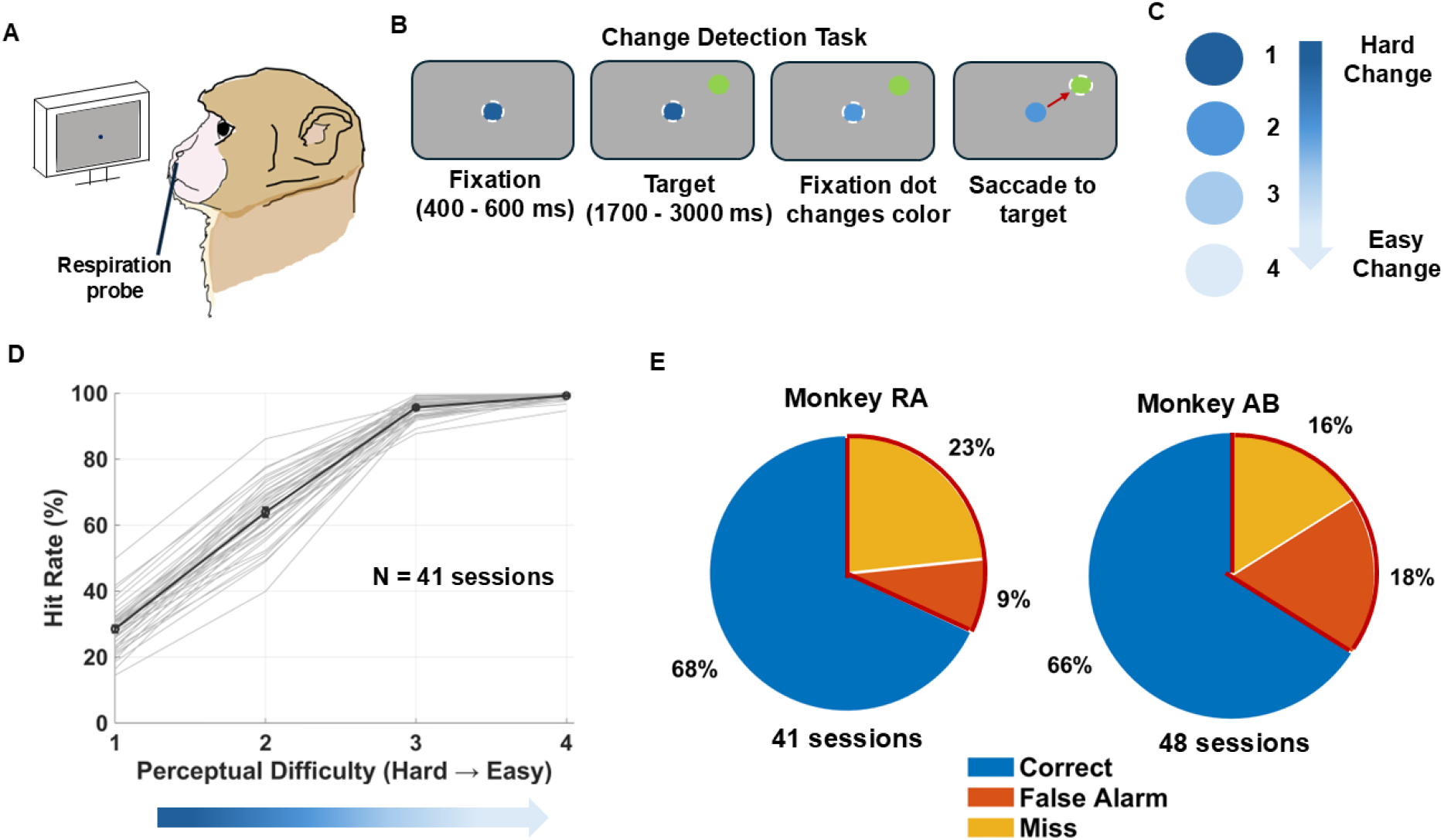
Respiration recording and behavioral performance during a visual perceptual task in nonhuman primates. (A)Experimental setup for simultaneous respiration recordings during perceptual change detection. (B)Schematic of the color change detection task, which involved a change in the color of the fixation spot quantified by a rotation in CIELUV color space (Monkey RA: ∼3°, ∼5°, ∼9°, ∼15°; Monkey AB: ∼4°, ∼8°, ∼10°, ∼12°). (C)Manipulation of perceptual difficulty by varying the degree of color rotation between fixation and target; larger rotations were easier to detect and smaller rotations more difficult. (D)Psychometric curve of Monkey RA’s behavioral performance. Hit rate (the number of correct trials divided by the total number of correct and miss trials) is plotted as a function of color-change difficulty (ranked from 1, the easiest to 4, the hardest). Light gray lines represent individual session performance, and the dark gray line shows the across-session mean (± SEM). (E)Behavioral performance summary across all sessions for each monkey. Pie charts indicate the proportion of correct (blue), false alarm (orange), and miss (yellow) trials for Monkey R (left) and Monkey A (right).

We analyzed each animal’s behavioral performance as a function of the color change magnitude, creating a psychophysical response function. Detection performance was defined as hit rate, calculated as the proportion of correct trials divided by the total number of correct and miss trials. This psychometric curve demonstrated a clear improvement in detection performance with increasing color change magnitude for individual sessions and the across-session average (light and dark gray curves, respectively, in Figure 1D for Monkey RA). We titrated the change difficulty values such that overall performance was similar for our two subjects, with approximately 70% of trials resulting in a correct detection (Figure 1E). Thus, our task structure created a robust framework for linking respiration with trial-by-trial behavioral outcomes.

### Session-level analysis revealed consistent differences in respiration timing based on trial outcome

To extract the features of respiration traces, we applied a multi-step signal processing pipeline (detailed description in STAR methods) and compared respiration waveforms across trial outcomes (Figure 2A–C). The overall frequency content of the respiration signal was relatively constant across the session, as seen in a spectrogram from one example session (Figure 2A). The dominant band of power was found between 0.1–0.5 Hz, around 15 breaths per minute. Example traces from Monkey RA (Figure 2B) illustrate how bandpass filtering with a third-order Butterworth filter (0.1–0.5 Hz) preserved the overall cycle structure while isolating the dominant frequency. These filtered respiration traces revealed trial outcome–specific differences in waveform shape. When aligned to trial start, average filtered traces across sessions (Figure 2C) showed subtle differences between correct and incorrect (false-alarm and miss) outcomes, motivating further feature-based analyses of respiration as a predictor of behavioral performance.

**Figure 2.**
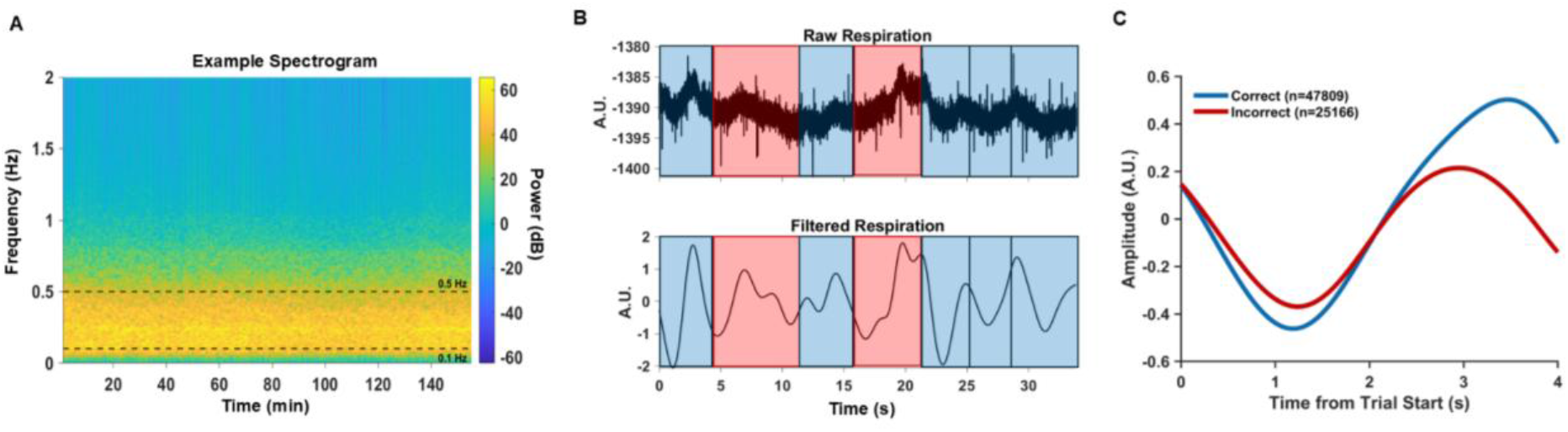
Signal processing pipeline and outcome-specific differences in respiration waveforms. (A)Respiration spectrogram from an example session. Power was concentrated between 0.1–0.5 Hz. (B)Example respiration traces from Monkey R. Top: raw respiration signal across several trials. Bottom: bandpass-filtered signal (0.1–0.5 Hz, third-order Butterworth filter). Blue shading marks correct trials, and red shading marks incorrect trials (false alarms + misses). (C)Average respiration waveforms aligned to trial onset, shown separately for correct (blue) and incorrect (red) trials. Shaded regions denote ± 95% confidence intervals across all sessions.

First, we wanted to analyze how respiration changes depending on trial outcome on a session-level basis. Our session-level analysis revealed consistent differences in respiration based on trial outcome. We wondered then whether specific components of the respiratory cycle related to behavior on individual trials. To do this we focused on the first full cycle after each trial’s onset and extracted a comprehensive set of temporal and amplitude-based features (Figure 3A). These included the onset times and durations of inhalation and exhalation, overall cycle length, and measures of volume and depth.

**Figure 3.**
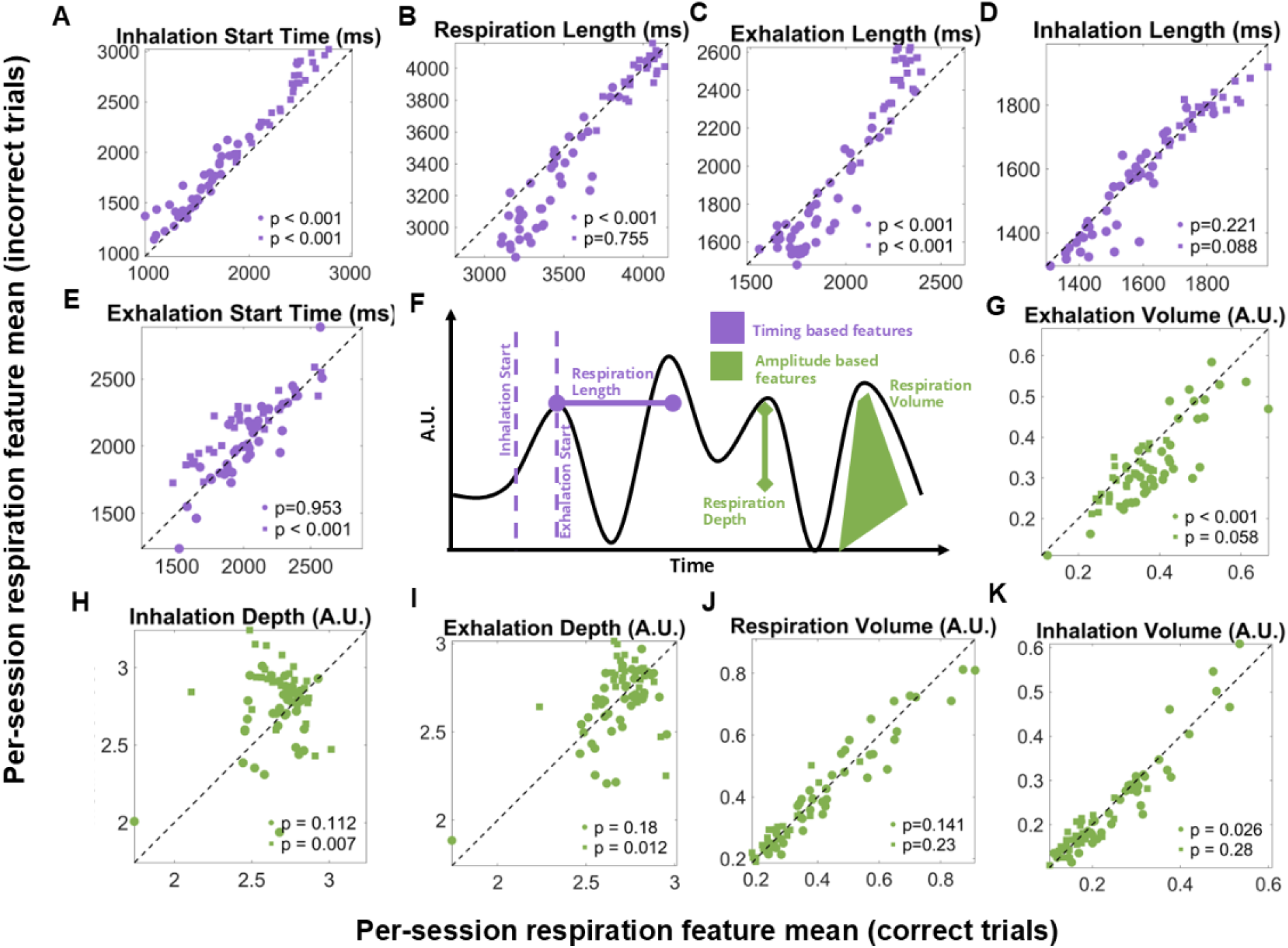
Session-level respiration timing features differ between correct and incorrect trials. (A–E) Session-level scatter plots of respiration timing-based features (purple), calculated as the mean feature value for correct trials vs. the mean for incorrect trials (y-axis). Each point corresponds to one session, with Monkey RA shown as circles (n = 41 sessions) and Monkey AB as squares (n = 48 sessions). Wilcoxon signed-rank test was used to determine significance. (F) Schematic illustration of respiration feature extraction. Timing features (purple) include inhalation/exhalation onset times, durations, and overall cycle length; amplitude features (green) include volume and depth. See STAR Methods for detailed definitions. (G–K) Same as (A-E) but grouped by respiration amplitude-based features (green).

Across sessions, timing features showed the strongest and most consistent outcome-related effects (Figure 3A–E). For example, inhalation onset relative to trial start (Figure 3A) was earlier for correct trials for both monkeys (Monkey RA: p < 0.001; Monkey AB: p< 0.001; pooled: p < 0.001). Respiration cycle length was also longer in correct trials for Monkey RA and the pooled data, though not for Monkey AB (Figure 3B), while exhalation length showed consistent increases across both animals (Figure 3C). In contrast, amplitude features did not show consistent effects across monkeys (Figure 3G-K). These effects point to a systematic alignment of the respiratory cycle with successful performance, suggesting that temporal aspects of breathing play a key role in shaping trial-by-trial behavior.

Importantly, not all timing features were the same between monkeys. Exhalation onset, for example, was significantly shifted in Monkey AB but not Monkey RA (Figure 3E), suggesting some degree of individual variability. Nevertheless, the overall pattern highlights that respiration timing reliably distinguished correct from incorrect outcomes at the session level. This motivated the next step of our analysis: moving beyond average session effects to ask whether these temporal features could predict behavior on a single-trial basis.

### Session-level distributions reveal that respiration features are discriminable within individual recording days

To complement our across-session comparisons, we next asked whether respiration features could differentiate between trial outcomes within a single recording session. Specifically, we examined whether respiration features that differed across sessions between correct and incorrect trials also showed separable trial-by-trial distributions within an individual day of recording. We focused on two respiration features that consistently showed outcome-related differences across sessions: respiration cycle length and exhalation volume. For illustration, we examined example sessions from the dataset shown in Figure 3. In one session, the distribution of respiration cycle length differed between outcomes, with correct trials tending to have longer respiration cycles than incorrect trials (Figure 4A). In another session, the distribution of exhalation volume was shifted such that correct trials tended to have larger exhalation volumes than incorrect trials (Figure 4B).

**Figure 4.**
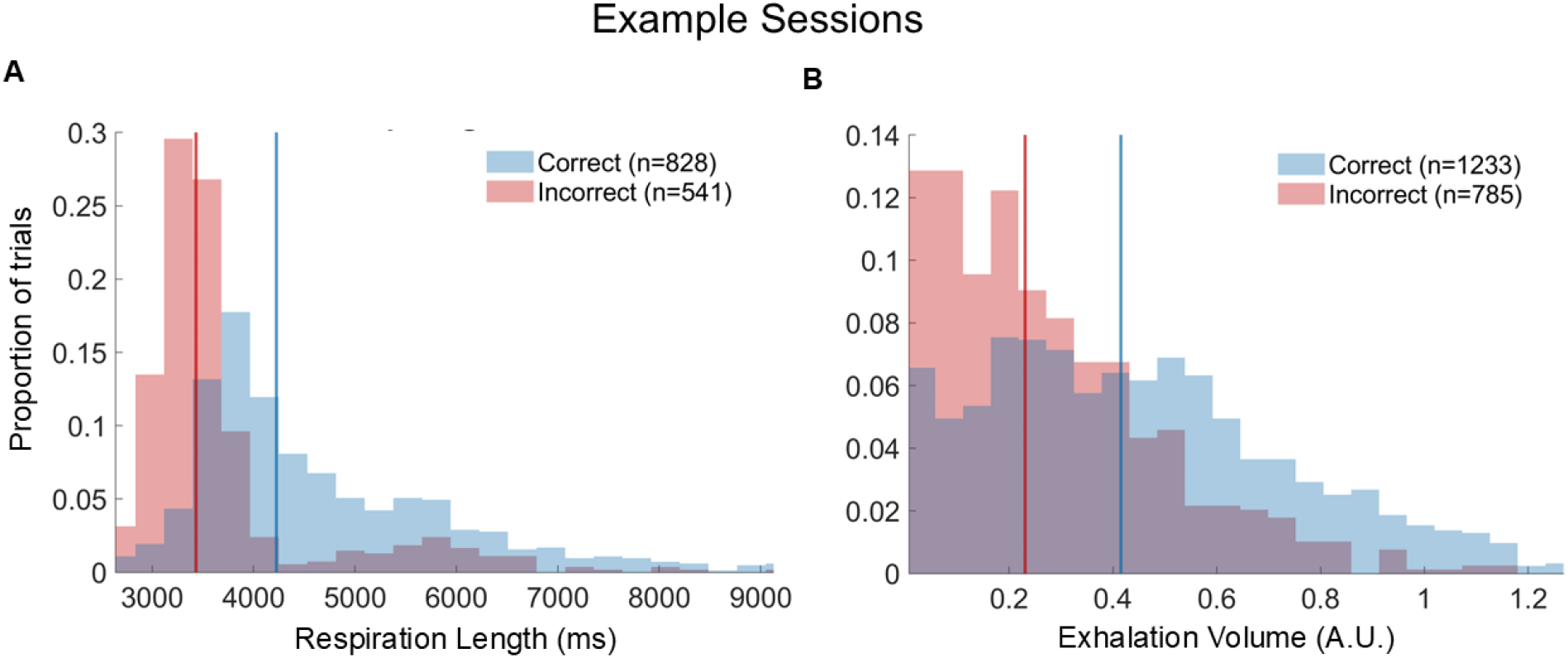
Respiration features are discriminable by trial outcome within an example session. (A)Example-session histograms of respiration length (first complete cycle after trial onset), plotted separately for correct (blue) and incorrect (red) trials. The mean of each distribution is indicated with a vertical line matched in color to the histogram. (B)Same as (A), but for exhalation volume from another example session. For both panels, distributions were compared using a two-sample t test, showing robust outcome-related differences (p ≪ 0.001 for each feature).

Although the trial-by-trial respiration values showed substantial overlap between outcomes, the distributions were significantly different within these example sessions (two-sample t-test; p ≪0.001 for both comparisons). These results demonstrate that respiration features carry information about behavioral outcomes that can be detected even within a single day of recording. We next turned to an approach that could aggregate across all of the respiration features to predict trial outcomes on a single-trial basis.

### Trial outcomes are well predicted by a combination of respiratory features

To test whether respiratory dynamics could predict behavior at the single-trial level, we trained a set of classifiers using features extracted from the first full respiration cycle following trial onset. Models included Gradient Boosting (GB), Random Forest (RF), Support Vector Machines (SVM), k-nearest neighbors (KNN) and multi-layer perceptrons (MLPs), providing a mix of interpretable ensemble methods and a neural network architecture.

Model performance was evaluated under two complementary cross-validation schemes (Table 1). We used 5-fold cross-validation (5CV) and leave-one-session-out (LOSO). The latter provided a particularly stringent test of robustness by assessing whether models trained on one set of recording days could generalize to entirely new sessions. Both validation schemes revealed that respiration features reliably predicted trial outcome. In Monkey RA, GB and MLP achieved the highest accuracy under 5CV (75%), while the GB performed best under LOSO (72%), closely followed by the MLP (71%). In Monkey AB, RF outperformed other models, reaching 73% in 5CV and 70% in LOSO. These results indicate that models can generalize both within and across sessions, confirming that respiration features carry behaviorally predictive information beyond session-specific variability.

**Table 1.** Cross-validated classifier accuracies for both monkeys.

| Model Type | Test Accuracy (%) |  |  |  |
| --- | --- | --- | --- | --- |
|  | 5CV -<br>Monkey RA | LOSO -<br>Monkey RA | 5CV -<br>Monkey AB | LOSO -<br>Monkey AB |
| Gradient Boosting | 75 | 72 | 72 | 70 |
| MLP | 75 | 71 | 72 | 70 |
| SVM | 73 | 70 | 71 | 67 |
| Random Forest | 73 | 70 | 73 | 70 |
| KNN | 69 | 67 | 69 | 65 |

Because respiration is a process that can span the border of trials, and potentially be impacted by recent events like sipping at the reward tube, we wanted to determine if the relationship between respiration and behavioral outcome could be impacted by trial history. To do this, we reran the GB classifier on a subset of trials which were chosen because they followed a correct trial. In doing this we balanced our trial selection by current outcome but holding preceding outcome constant. We found that classifier accuracy remained comparable to prediction on the full set of trials, reaching 74% in Monkey RA and 72% in Monkey AB. This demonstrates that respiration features continue to predict behavior even after controlling for trial history.

To interpret which aspects of respiration drove model performance, we examined feature importance scores from the GB model (Figure 5A–B). Timing features, such as respiration length and inhalation start time, were consistently more informative than amplitude-based features, aligning with results in Figure 3. We next addressed feature redundancy using an exhaustive feature-combination analysis (Figure 5C–D). For each possible number of features, all combinations were evaluated and test accuracy was summarized as the mean across all possible sets (gray line) and the best-performing set (red line). In both animals, accuracy increased with the addition of features and the best set of 3-6 features achieved performance on par with the full set. Timing features again were represented in these smaller feature combinations, with amplitude features providing modest complementary value as the model started to reach its plateau. We also built a model which pooled trials across both monkeys into a single dataset yielding comparable accuracy (72% with GB), which indicates that despite individual variability the predictive structure of respiration features generalized across animals.

**Figure 5.**
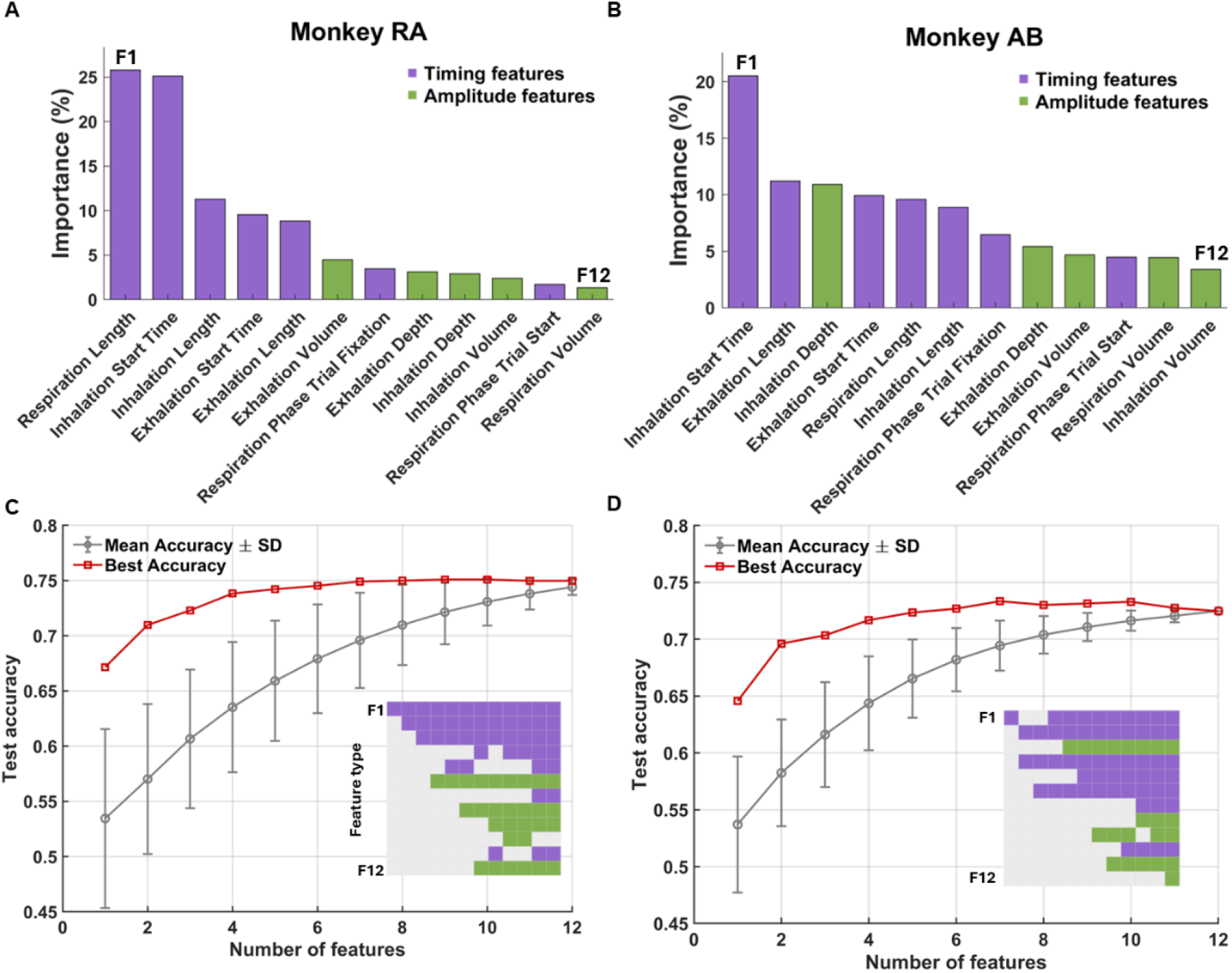
Features related to respiration timing were the strongest predictors of trial outcome. (A)Gradient Boosting feature importance scores for Monkey RA, with features color-coded by type (timing = purple, amplitude = green). (B)Same as (A), for Monkey AB. (C)Test accuracy as a function of the number of features included in the model for Monkey RA. For each feature count, all possible feature combinations were evaluated. The gray line shows mean accuracy across combinations, and the red line indicates the best-performing combination (5-fold CV). The inset colored grid below the curve depicts which features were present in the best-performing combination at each feature count: each square corresponds to a feature, that follows the same order as the feature importance from (A) and (B) (y-axis) and number of features in each iteration (x-axis), with purple denoting timing features and green denoting amplitude features. (D)Same as (C), for Monkey AB.

## DISCUSSION

Our findings demonstrate that respiration carries reliable predictive information about perceptual performance on a single-trial basis. By using interpretable, feature-based models, we identified timing-based respiratory features, particularly the latency to inhalation onset and the duration of the first cycle after trial onset, as the strongest predictors of trial outcome. In contrast, amplitude-based features such as depth or volume contributed far less. A key strength of our approach is the systematic extraction of cycle-level features (onset times, durations, phase, depth, and volume) aligned to task behavior.

Across trials, correct responses were associated with earlier inhalations and longer cycle durations, indicating that when the subject started breathing in the trial was behaviorally meaningful. This aligns closely with human behavioral work showing that the inhalation phase of breathing often synchronizes with task engagement and increases the probability of detecting near-threshold stimuli. For example, respiration phase at stimulus onset predicts visual detection performance^14^, and humans spontaneously time inhalations to anticipated cognitive demands^15,39^. Although most prior studies describe effects in terms of phase, our results show that explicit timing (i.e., when inhalation begins relative to trial onset) provides an equally powerful description of how respiration shapes perception. In mice, combinations of temporal features such as inspiratory and expiratory durations reliably differentiate behavioral states over extended timescales^27^. Our results extend this logic to moment-to-moment perceptual decisions, demonstrating that fine-grained temporal features of a single respiration can predict whether a stimulus will be detected.

Together, these behavioral findings complement a growing neural literature showing that respiration rhythmically modulates cortical excitability. Human EEG/MEG studies reveal respiration-locked fluctuations in alpha and beta power, with phases of heightened excitability corresponding to improved stimulus detection^14,15^. Such oscillatory modulations suggest a form of sensory gating, where inhalation-aligned increases in cortical activity make the brain more receptive to incoming sensory information. This framework provides a mechanistic interpretation of our behavioral results: trials that begin during “high-excitability” respiratory phases, marked by earlier inhalation onset or longer initial cycles, may benefit from increased neural excitability and therefore show higher perceptual accuracy. This work supports a hypothesis in which respiration shapes trial-by-trial fluctuations in perceptual performance. A natural next step is to add neural recordings to directly test whether these behaviorally advantageous respiratory mechanisms correspond to measurable changes in cortical excitability and sensory processing.

This strong contribution of timing-based respiratory features to predicting trial outcomes aligns well with broader evidence that systemic physiology is connected to brain activity and behavior. In ECG and pupil studies, trial onset is often accompanied by reliable shifts in heart rate and pupil size, reflecting preparatory autonomic adjustments^16,40–43^. Similarly, work in monkeys and humans have shown that neural activity in the visual cortex modulates prior to stimulus onset in anticipation of task demands^40–43^. These findings suggest the existence of latent preparatory states that coordinate interoceptive and neural rhythms to optimize sensory processing.

More broadly, theories of interoception suggest that bodily rhythms provide context signals that dynamically influence cognition^44,45^. One account proposes a common arousal mechanism, in which brainstem nuclei such as the locus coeruleus coordinate global state changes that simultaneously modulate cortical excitability and peripheral physiology, including respiration^46,47^. A complementary framework posits a direct respiratory-neural coupling, where respiratory rhythm generators and afferent pathways interact with cortical and thalamic circuits, allowing breathing dynamics themselves to entrain neuronal oscillations and shape sensory processing^48,49^. Our data provides complementary evidence that respiration participates in this preparatory alignment and may improve detection accuracy. One interpretation is that breathing rhythms help structure the timing of cortical excitability, such that trials aligned to favorable phases of the respiratory cycle are more likely to succeed. Future work should test this causal role directly, for example, by manipulating respiration timing relative to task events and assessing the impact on performance.

Several considerations point to clear directions for future work. First, while our classifiers exceeded chance and generalized across sessions, they likely capture only a portion of the relevant variance, leaving room to test whether combining respiration with complementary signals (e.g., pupil, ECG, LFP/EEG) and richer temporal models improves predictive power and mechanistic insight^31,32,50^. Second, recording respiration is indirect and often relies on secondary measures^51^. In our case, the temperature in the nose was a high-resolution measure for respiration but lacked an absolute amplitude calibration. Future designs can incorporate stable calibration references to better capture amplitude effects. Third, we studied two animals performing a single task. An advantage of our paradigm in non-human primates is that we collected highly replicable behavior in well trained subjects across thousands of trials and many sessions. Nonetheless, expanding to larger, mixed-sex cohorts and multiple task contexts will test generalizability and boundary conditions.

To summarize, this study shows that respiration is not merely a background rhythm but a dynamic indicator of behavior. By revealing that fine-grained features of respiration, especially its task-related timing, predict perceptual performance at the single-trial level, we emphasize the need to incorporate interoceptive rhythms into models of cognition. This framework opens the door to testing whether respiration-aligned interventions could enhance perception, attention, or learning in health and disease.

## RESOURCE AVAILABILITY

### Lead contact

Further information and requests for resources should be directed to and will be fulfilled by the lead contact, Matthew A. Smith.

### Materials availability

This study did not generate new materials.

### Data and code availability

Data and analysis code used in this paper will be available as of the date of publication.

## ACKNOWLEDGMENTS

The authors would like to thank our animal care staff for their support. This work was supported by NIH R01 MH128393. E.S. was supported by NIH T32 EB029365 and the Ronald F. and Janice A. Zollo Fellowship. D.I. was supported by NIH F30 MH129056, the Ronald F. and Janice A. Zollo Fellowship, the Thomas-Pittsburgh Chapter ARCS Award and the Carnegie Mellon University Neuroscience Institute Carnegie Fellowship.

## AUTHOR CONTRIBUTIONS

Conceptualization, E.S., J.M.K and M.A.S.; methodology, E.S., D.I., M.G., S.N., J.M.K and M.A.S Investigation; E.S., D.I., M.G. and S.N.; writing—original draft, E.S.; writing—review & editing, E.S., D.I., M.G., S.N., J.M.K. and M.A.S.; funding acquisition, J.M.K and M.A.S.; resources, J.M.K. and M.A.S.; supervision, J.M.K. and M.A.S.

## DECLARATION OF GENERATIVE AI AND AI-ASSISTED TECHNOLOGIES

During the preparation of this work, the author(s) used ChatGPT in order to assist with wording and editing of the writing. After using this tool, the author(s) reviewed and edited the content as needed and take full responsibility for the content of the publication.

## STAR?METHODS

### KEY RESOURCES TABLE

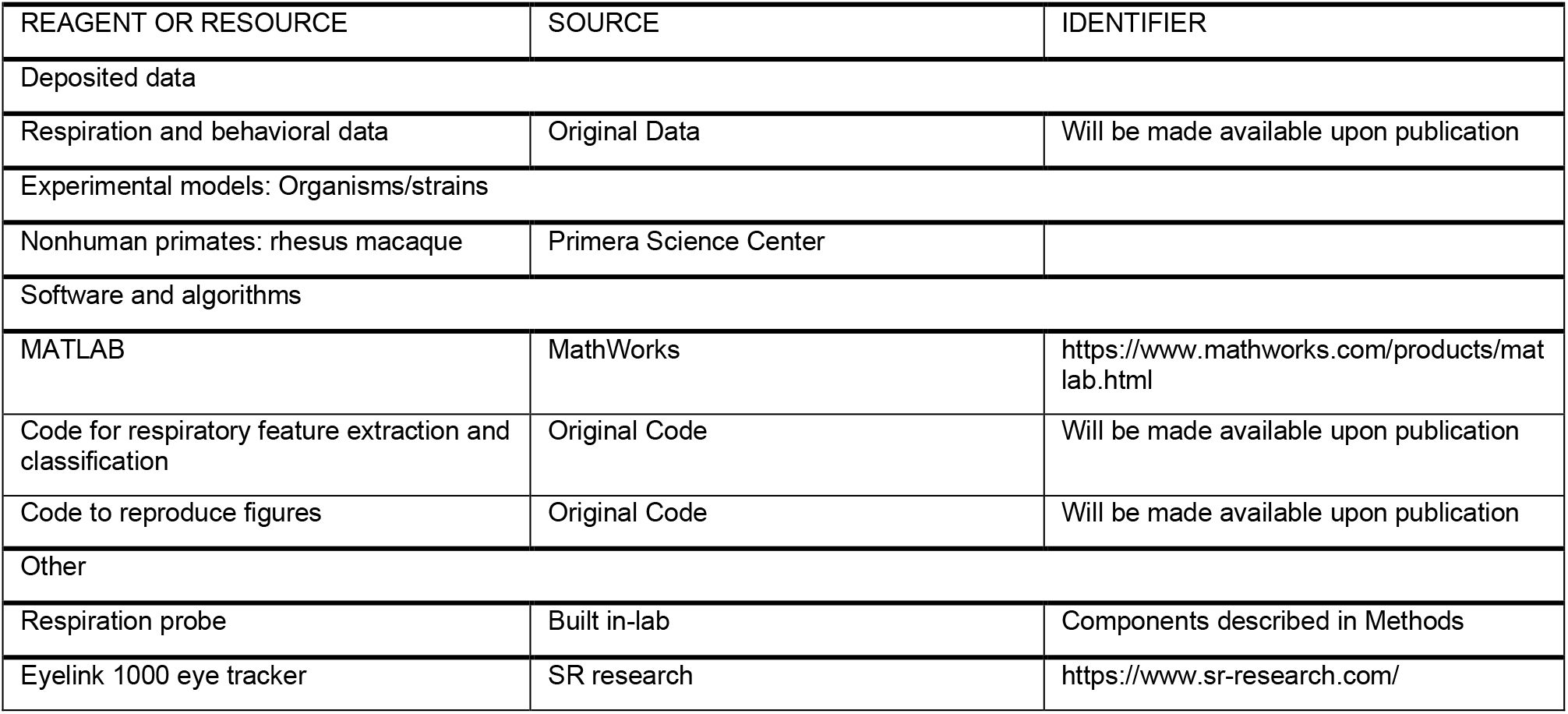

### EXPERIMENTAL MODEL

Two male rhesus macaque monkeys (Macaca mulatta) were used in this study. They were 10 (Monkey RA), and 9 (Monkey AB) years old at the time of data collection. The animals were housed singly in a room operating on a standard 12 hour light/dark cycle and were provided with an enhanced enrichment program. All experimental procedures were approved by the Institutional Animal Care and Use Committees of Carnegie Mellon University and were in accordance with the United States National Research Council’s Guide for the Care and Use of Laboratory Animals.

### METHOD DETAILS

#### Respiration recordings

Respiration was recorded continuously using a custom-built nasal thermocouple probe positioned at the entrance of one nostril in each monkey. The probe consisted of a Type-K thermocouple (glass braid insulated) connected to an AD8495 thermocouple amplifier (Adafruit). The thermocouple wire was threaded through a thin plastic rod and mounted on a cable tension based flexible arm to allow precise positioning in front of the nostril without contacting the animal.

Respiration was measured via temperature fluctuations in the airflow from the nostril. Exhalation caused an increase in temperature relative to the ambient room air (∼70° F) and then inhalation then caused a relative decrease in temperature as the room air was drawn past the thermocouple probe, producing corresponding voltage changes at the amplifier output (increase in voltage for and increase in temperature). The analog signal from the thermocouple amplifier was routed through a BNC connection to the data recording system (Ripple Inc; Salt Lake City, UT) and acquired using Trellis software on an analog channel sampled at 1 kHz.

#### Eye Tracking

Eye position was recorded using an infrared eye tracking system at a rate of 1000 Hz (EyeLink 1000; SR research).

#### Behavioral task

Subjects performed a color-change detection task in which their goal was to detect and respond as quickly as possible to a subtle change in the color of a central fixation dot. Each trial began with a blank screen presented for a brief inter-trial interval (trial start), after which a blue fixation point appeared at the center of the screen. The monkey initiated the trial by directing its gaze to this fixation point (fixation onset; 321 ± 57 ms from trial start) and was required to maintain fixation for an initial period of 400–600 ms (chosen randomly).

After this initial fixation period, a peripheral target appeared at one of eight possible locations spaced 45° apart around the fixation point. Unlike a simple visually guided saccade task, the animal was required to continue fixating through an additional delay period while the target remained present. The duration of this delay was drawn from a truncated exponential distribution (mean: ∼2000 ms, range: 1700–3000 ms), preventing temporal predictability of the go cue.

At the end of the delay, the central fixation dot changed color. This color rotation in CIELUV color space served as the go cue, indicating that the monkey should look toward the previously presented peripheral target. The magnitude of the color rotation varied across trials (Monkey RA: by ∼3°, ∼5°, ∼9°, ∼15°; Monkey AB: by ∼4°, ∼8°, ∼10°, ∼12°) with larger rotations being perceptually easier to detect.

A trial was counted as correct if the monkey made a saccade to the target within the allowed reaction time window (<500ms to start a saccade and, <200ms to complete a saccade). Trials were labeled false alarms if the monkey broke fixation and looked toward the target before the color change occurred and misses if the monkey failed to initiate a saccade during the reaction time window after the color change. Successful trials earned a fixed juice reward.

Following trial completion, the timing of the subsequent trial depended on the outcome of the preceding trial. After correct trials, the next trial began on average 0.99 ± 0.06 s after the saccade to the target (this interval included both an enforced wait period and the time required for the experimental control system to prepare the next trial). Following false alarms, a timeout penalty was imposed, resulting in a longer inter-trial interval; the next trial began 1.62 ± 0.05 s after the saccade. After miss trials, in which no saccade was made to the target, a longer timeout period was enforced before the next trial began, resulting in an average interval of 2.00 ± 0.10 s after the color change.

### Data Analysis

#### Respiration waveform feature extraction

The respiration signal was processed using a third-order Butterworth bandpass filter (0.1–0.5 Hz) to isolate the typical frequency range of respiration. To eliminate residual high-frequency noise, additional notch filters were applied using the “iirnotch” function (MATLAB 2024b, MathWorks Inc.). Following filtering, the signal was inverted so that rising phases correspond to inhalation and falling phases to exhalation. Finally, the signal was z-scored to account for potential variability in probe placement across recording sessions and then epoched into trials.

A range of respiration features were extracted using different signal processing methods. For each trial, inhalation and exhalation phases were identified using a tailored implementation of the “findpeaks” function (MATLAB 2024b, MathWorks Inc.). Peaks (marking the start of exhalation) and valleys (marking the start of inhalation) were detected to determine the timing and structure of each respiratory cycle. From these, respiration length (interval between successive peaks), as well as inhalation and exhalation durations (intervals between peaks and valleys, and vice versa), were calculated. The onset of inhalation and exhalation were measured relative to trial start to assess timing relative to the task. Additionally, the prominence of each detected peak was extracted to identify for inhalation/exhalation depth. Finally, the respiration signal was numerically integrated between inhalation and exhalation boundaries to compute relative inhalation and exhalation volumes, as well as total respiration volume.

Respiration phase was established by applying the Hilbert transform on the filtered, z-scored respiration signal of the whole session. Then the “angle” function (MATLAB 2024b, MathWorks Inc.) was applied to understand how phase (in degrees) changes across the session. Finally, the respiration phase in degrees was computed separately relative to the trial start and fixation onset.

#### Data preparation

For each trial, a set of respiratory features was extracted from the first complete respiration cycle following trial start. This cycle was defined as the interval from the first inhalation onset to the subsequent inhalation onset. Focusing on this initial cycle allowed for consistent feature extraction across trials and ensured that all features were temporally aligned relative to the behavioral task.

Each trial was represented as a row in a feature matrix of size [number of trials × number of features], with the features mentioned above.

Before modeling, each feature column was z-score normalized across all to place all features on a common scale. The finalized design matrix and label vector were then used as inputs for all machine learning classifiers.

#### Machine Learning classification

Data from 41 recording sessions from Monkey RA and 48 from Monkey AB of respiration waveforms were used to predict behavioral outcomes based on features extracted from the respiratory signal. Two validation strategies were implemented to evaluate model performance: (1) five-fold cross-validation, with 80% of trials used for hyperparameter tuning and training and 20% for testing, and (2) leave-one-session-out (LOSO) validation, where models were trained on all sessions except one, which was held out for testing. Both approaches ensured that evaluation was robust across trials and sessions, helping to minimize overfitting and assess generalizability across animals and experimental conditions.

All models were implemented in MATLAB. The classifiers tested were Random Forrest, Support Vector Machines, k-Nearest Neighbor Search, a boosted random forest, ensemble method using decision trees, and multi-layer perceptron (MLP). Model performance was quantified using the area under the curve (AUC), confusion matrices, and accuracy (%).

### QUANTIFICATION AND STATISTICAL ANALYSIS

We report *n* as the number of experimental sessions per animal (Monkey R: 41; Monkey A: 48), each on a separate day. Session-level differences in respiration features between correct and incorrect outcomes were assessed with two-sided Wilcoxon signed-rank tests *(α* =0.05) and with two-sample t-tests *(α* =0.05). For single-trial decoding, model performance is summarized by classification accuracy (%).

All analyses were implemented in MATLAB R2024b (MathWorks; Statistics and Machine Learning Toolbox); circular tests used standard implementations (e.g., Circular Statistics Toolbox). Our internal lab software randomized the trial conditions; analyses were automated and performed blind to session identity. Sessions/trials failing predefined quality-control thresholds (e.g., corrupted respiration or insufficient incorrect trials for comparison) were excluded prior to analysis, with criteria and totals detailed in figure legends.

